# A group refactorization procedure for sleep electroencephalography

**DOI:** 10.1101/2020.12.29.424768

**Authors:** Robert G. Law, Shaun M. Purcell

## Abstract

[WORKING DRAFT] In order to relate health and disease to brain state, patterns of activity in the brain must be phenotyped. In this regard, polysomnography datasets present both an opportunity and a challenge, as although sleep data are extensive and multidimensional, features of the sleep EEG are known to correlate with clinical outcomes. Machine learning methods for rank reduction are attractive means for bringing the phenotyping problem to a manageable size. The whole-night power spectrogram is nonnegative, and so applying nonnegative matrix factorization (NMF) to separate spectrograms into time and frequency factors is a natural choice for dimension reduction. However, NMF converges differently depending on initial conditions, and there is no guarantee that factors obtained from one individual will be comparable with those from another, hampering inter-individual analysis.

We therefore reseed time-frequency NMF with group frequency factors obtained from the entire sample. This “refactorization” extends classical frequency bands to frequency factors. The group reseeding procedure coerces factors into equivalence classes, making them comparable across individuals. By comparing frequency factor properties, we illustrate age-related effects on the sleep EEG. The procedure can presumably be adapted to higher resolutions, e.g. to local field potential datasets, for characterizing individual time-frequency events.

## 1 Introduction

With the advent of large-scale neural time-series datasets, an open question is how best to distill these data into quantitative phenotypes. Nowhere is this more pertinent than in polysomnography, where a single recording session includes upwards of eight hours of electroencephalography (EEG) as well as a variety of other physiological time series. A common reduction of data in sleep EEG is to calculate the power in a small set of frequency bands for each epoch, and then examine the dynamics of bandlimited power over the course of the night [2]. Power in each of these bands typically corresponds to oscillatory intensity, number, and/or extent and are markers for a variety of healthy and diseased states (e.g. [1, 3, 20, 13]). Yet it is well-known that the frequency modes of comparable oscillations vary across individuals and with age (e.g. [3]), and despite nearly a century of research there remains disagreement on even the boundaries of each frequency band (compare, for instance, [15] to [7]).

In small datasets such definitional issues can be dealt with heuristically, but in large datasets they may be more problematic: Directly comparing 20Hz activity across a population is suboptimal, if 20Hz activity in one individual is functionally equivalent to 18Hz activity in another. Furthermore, the observed activity at a specific frequency bin (e.g. 15 Hz) or band (e.g. sigma) may reflect a summation across a heterogeneous mixture of components, which may individually exhibit qualitatively distinct patterns of association, or dynamics, whereas analyses based only on the observed spectrogram may obscure such features.

Our contribution here is to establish a data reduction procedure based on nonnegative matrix factorization (NMF; [14]) that constructs individualized frequency factors that can be meaningfully interpreted and compared at the population level, and which potentially deconvolves distinct but overlapping spectral signatures, and to show that these factors vary with age in an interpretable way.

## 2 Methods

### 2.1 Datasets

We studied two datasets from the National Sleep Research Resource polysomnography database [22]. The Childhood Adenotonsillectomy Trial (ChAT) was chosen for future examination of test/retest correlations ([17]; *n* = 859 recording sessions; 452 baseline, 407 followup; aged 5–10 years), while the Cleveland Family Study (CFS) was chosen as a dataset for investigating age correlations here ([18]; *n* = 540 individuals; aged 7–89 years; see below).

### 2.2 Preprocessing

All preprocessing was performed using Luna, a freely available command-line utility specialized for handling large-scale sleep data (http://zzz.bwh.harvard.edu/luna). Each EEG time-series (channel C3; contralateral mastoid reference) was converted to microvolts and partitioned into nonoverlapping, contiguous 30-second epochs. A spectrum for each epoch was estimated using the Welch method (0.5 − 45*Hz*; 0.5*Hz* bins). For each time-series with *m* epochs, this yields a non-overlapping spectrogram **Y**, a 90 × *m* matrix representing the time-frequency plane. Further analysis was restricted to sleep and brief arousals within sleep cycles [6].

For polysomnograms in the Cleveland Family Study, we removed all sessions where sleep onset time was before 10pm, as participants were awoken at this time for clinical assessment. Due mainly to the denoising nature of NMF [21], we found that no further artifact rejection was necessary for our purposes here.

### 2.3 Factorization

Nonnegative matrix factorization [14] finds a fixed-rank approximation **WH ≈ Y** with all entries nonnegative, enabling a “parts-based” decomposition [10] in the sense that each epoch of spectral data is reconstructed by combining a small number of spectral ingredients **w** (“what”) in various amounts **h** (“how much”). This corresponds to the notion that a spectrum is generated by a finite collection of processes, each with its own spectrum, that are combined at the scalp-level. Unlike global methods like principal components analysis, the approximation reached by NMF depends critically on the initial seed matrices **W**_0_, **H**_0_. While many applications of nonnegative matrix factorization use random seeds [4], “warm-start” factorization methods [5] can outperform random seeds, although they rely on the approximate correctness of an initial guess.

Here, we construct such an initial guess by sampling from the population. Spectrograms were first log-transformed and minimized **Y**′ ← log(^**Y**^/_min **Y1**_), so that min **Y**′ = 0. All factorizations were then performed using the multiplicative method [10] implemented in the NMF R package [8]; see Figure 1 for an example.

**Figure 1:**
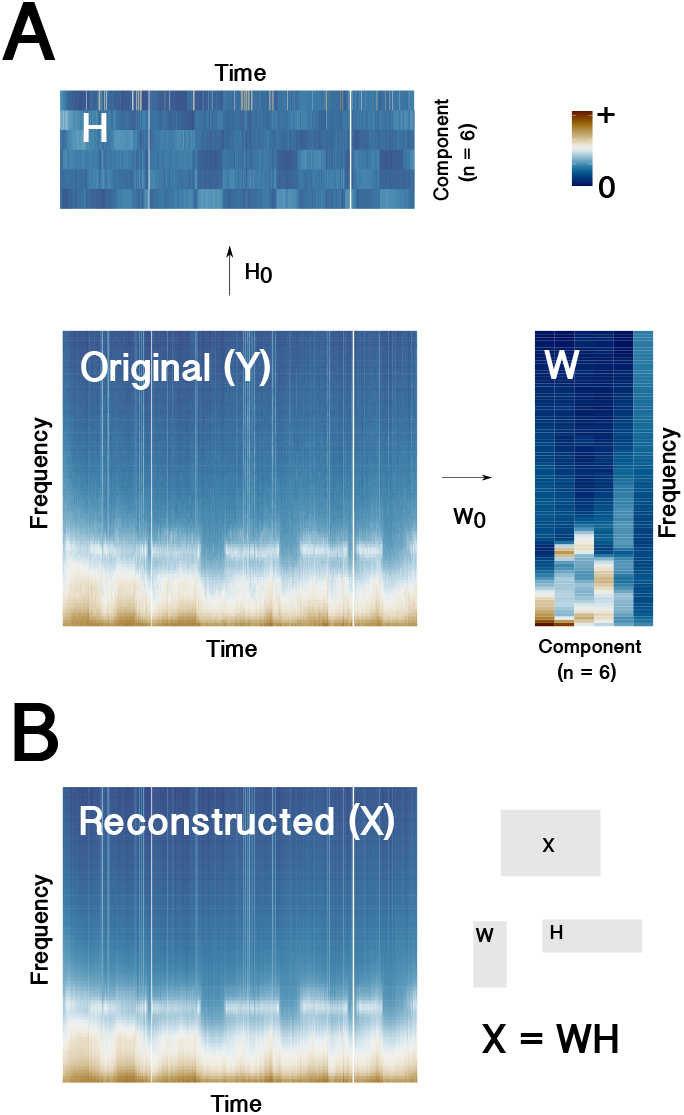
**A)** Factorization of a log-spectrogram **Y** into nonnegative frequency- and time-components **W** and **H**. The initial conditions for each factorization are shown as arrows. **B)** Approximate spectrogram, reconstructed by multiplication of the factors.

We take a three-step approach (Figure 2): We first presuppose a nearly-uniform seed, where **H**_0_ is uniform (*h_ij_* = 1) and **W**_0_ differs from uniform only enough to break symmetry in the initial conditions (*w_i,i_* = 2; *W_ij_* = 1 otherwise). This yields initial factorizations for each individual *k*: **W**_*k*_, **H**_*k*_. We discard the temporal factors **H**, and then concatenate the frequency factors from this first factorization, as **W**_1_|**W**_2_| … |**W**_*k*_| … |**W**_*n*_. We factorize this concatenated matrix to obtain six group-level frequency factors in a 90 × 6 matrix **Ω**. These are used as frequency seeds (again with uniform temporal seeds **H**_0_) for a second NMF round on individuals, from which the final frequency and temporal factors are obtained.

**Figure 2:**
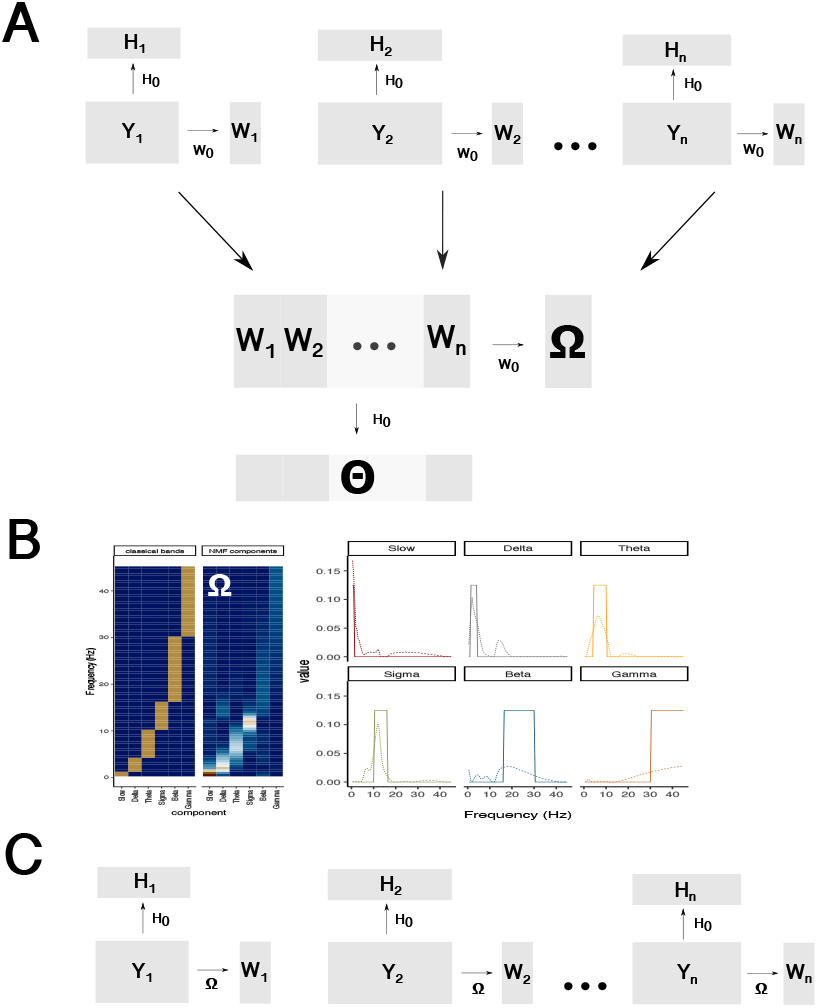
**A)** The **W**_*k*_ obtained from a first factorization using nearly-uniform seeds **W**_0_, **H**_0_ (see text) are concatenated and themselves factored to obtain a set of group factors **Ω**. **B)** Comparison of classical frequency bands to Ω-factors as heatmaps (*left*) and as functions (*right*) **C) Ω**-factors are then used as seeds to obtain a final factorization.

More informally, we should like to: 1) use NMF with a nearly-uniform seed to find six spectral ingredients in each individual’s sleep 2) Put all 6 × n individual ingredients together and use NMF to find six “base ingredients”, and then 3) Look for these base ingredients (and variations thereof) by seeding a second NMF round with the base ingredients. We examine time and frequency factors obtained from this second round of factorization.

Note that when we simply used the factors obtained from (1), the factorization was not stable: i.e. similar factors were permuted across sessions, making comparisons of factors across sessions effectively impossible. Evidently, though, the group seed **Ω** contains zero entries unique for each factor (Figure 2B), all of which are preserved by the NMF update rule. As such, each factor has a unique support that is preserved in each iteration of NMF: This ultimately defines an equivalence class of probability density functions (e.g. normalized spectra) admissible for each factor, which is what makes within-class comparisons reasonable.

## 3 Results

### 3.1 NMF frequency factors extend classical frequency bands

Inspection of the frequency factors then yielded an immediate visual match to classical frequency bands in both the ChAT and CFS datasets (Figure 3A/B), although the classical theta and alpha bands merged into one factor (denoted “*theta*” henceforth). The primary difference between classical bands and NMF factors is again one of support: the **w**-factors – while concentrated within classical bandlimits – are also supported outside those limits. Individual **w**-factors clustered near their respective means. Factors were comparable between the two datasets (Figure 3C), but all aside from the “gamma” factor were distinguishable between ChAT and CFS, with presumably age-dependent shifts evident in “delta”, “theta”, “sigma”, and “beta” factors.

**Figure 3:**
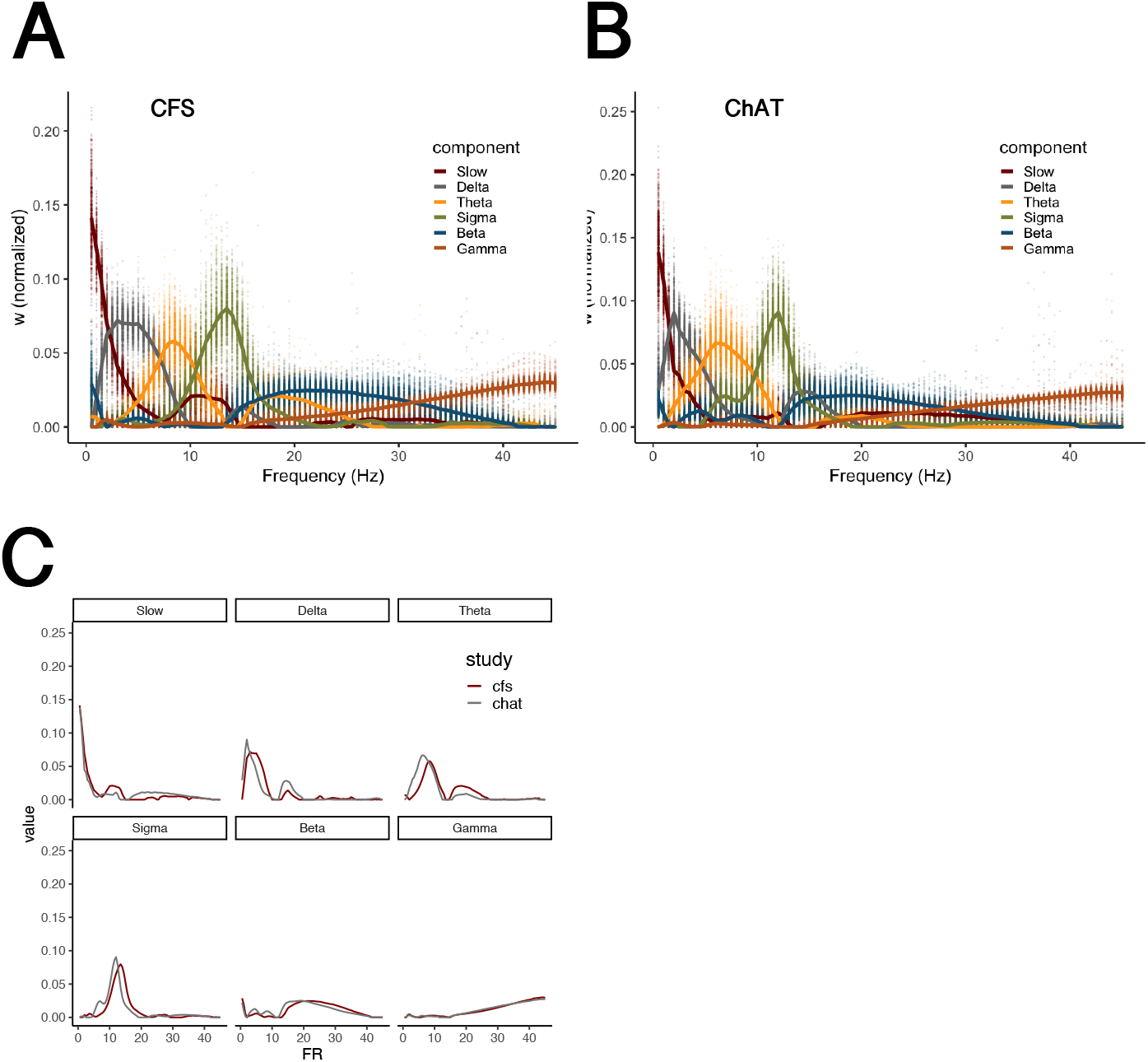
Frequency factors in the Childhood Adenotonsillectomy Trial and Cleveland Family Study. **A)** ChAT frequency factors. Mean and all individual data points are shown. *Inset*: Strength of traitlikeness is indicated by the z- score of the actual test/retest correlation relative to a distribution of correlations under shuffled session IDs. **B)** CFS frequency factors. **C)** *Left*: example of frequency factors with good test/retest correlation. *Right*: example with poor correlation. **D)** Overlaid mean frequency factors from the ChAT and CFS datasets.

### 3.2 Frequency factors reveal age-dependent cross-frequency coupling

We next considered how frequency factors were correlated with age – a well-studied demographic covariate of sleep dynamics (Figure 4). We found that age differences within factors tended to occur when spectral power can be grouped preferentially into either one factor or another at the population level. For instance, since the “delta” factor and the “sigma” factor have support near 15Hz, at each epoch NMF can couple each unit of 15Hz power to either 1–4Hz or ~12Hz power. We find a strong age-dependence to this assignment, with aging associated with less assignment of 15Hz power to the “delta” factor, reflecting reduced coupling of 15Hz power to the 1–4Hz delta mode with age (Figure 4A). With increasing age, 15Hz power is instead assigned to the “sigma” factor (Figure 4B). In other words, fast spindles appear to decouple from slow oscillations with age, agreeing with results from a recent slow-wave stimulation experiment [19]. Coupling of 10Hz power to either the “theta” or “slow” factor was also age-dependent (Figure 4C/D).

**Figure 4:**
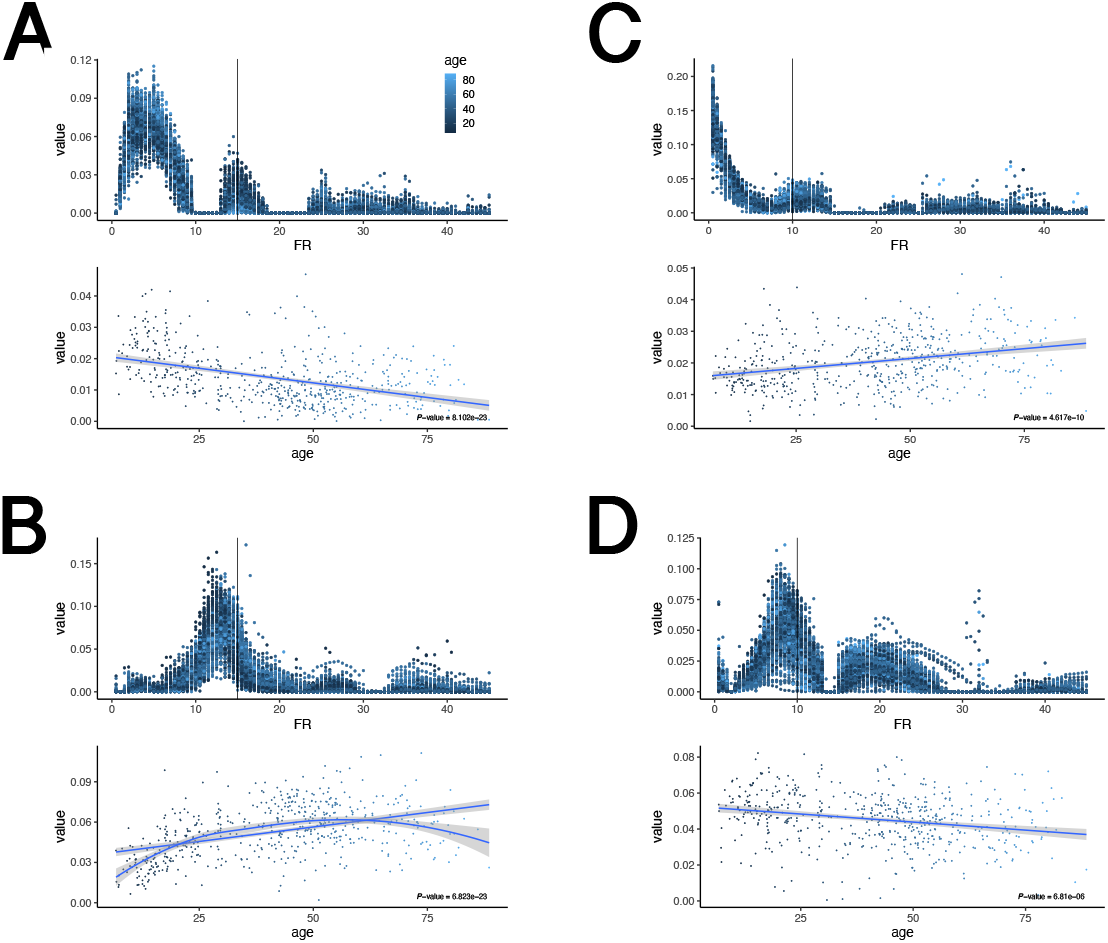
Frequency factor correlates with age. **A)** *Top*: Individual “Delta” frequency factors are plotted, colored by age. The vertical line represents a slice through the frequency of interest, revealing (*bottom*) an age-anticorrelated 15Hz component. **B)** “Sigma” factor has a corresponding age-correlated 15Hz component. **C)** “Slow” factor has an age-correlated 10Hz component. **D)** “Theta” factor has a corresponding age-anticorrelated 10Hz component. All p-values are uncorrected (540 comparisons).

## 4 Discussion

We have developed a procedure for extracting comparable EEG phenotypes in sleep using nonnegative matrix factorization, showing that group refactorization successfully decomposes power spectra into frequency factors resembling but extending classical frequency bands, with age-dependent features. While our efforts here were primarily concerned with validating the procedure, applications may range from cross-species comparisons of factors to subepoch characterization of individual time-frequency events such as slow waves and sleep spindles, along the lines of [16].

Nonnegative methods have been applied to polysomnography previously: The most similar approach to ours is [9], who used nonnegative tensor factorization with twelve fixed frequency bands to discriminate spectrotemporal patterns in Alzheimer’s disease, but this factorization was supervised. Tensor factorization, moreover, cannot be applied to sleep recordings with variable duration without truncation; this indeed was a primary motivation in developing our method. Other related approaches are [11] and [12]: [11], in particular, applied kernel NMF to obtain frequency factors within individuals; these factors, however, were not compared across individuals as the primary motivation for the study was brain-computer interface design.

It is interesting that we were able to obtain our results with a standard NMF algorithm using only manipulations of the seed matrices: It is not completely clear why the uniform seed **H**_0_ appears to yield sufficient smoothness for our purposes, nor why the group seed **Ω** is sparse enough to assure equivalance of factors across individuals. However, given these findings, we are left with the possibility that in combination with experiment and modeling, phenotypic factors obtained from these epidemiological-scale datasets might be mapped to physiological properties of cells or networks, in a more direct manner than using classical frequency bands.

